# Quantifying viral pandemic potential from experimental transmission studies

**DOI:** 10.1101/2025.03.24.645081

**Authors:** Elizabeth D. Somsen, Kayla M. Septer, Cassandra J. Field, Devanshi R. Patel, Anice C. Lowen, Troy C. Sutton, Katia Koelle

## Abstract

In an effort to avert future pandemics, surveillance studies aimed at identifying zoonotic viruses at high risk of spilling over into humans act to monitor the ‘viral chatter’ at the animal-human interface. These studies are hampered, however, by the diversity of zoonotic viruses and the limited tools available to assess pandemic risk. Methods currently in use include the characterization of candidate viruses using in vitro laboratory assays and experimental transmission studies in animal models. However, transmission experiments yield relatively low-resolution outputs that are not immediately translatable to projections of viral dynamics at the level of a host population. To address this gap, we present an analytical framework to extend the use of measurements from experimental transmission studies to generate more quantitative risk assessments. Specifically, we use within-host viral titer data from index and contact animals to estimate parameters relevant to transmission, including an estimate of transmissibility. We then extended this model to estimate epidemiological parameters, such as the basic reproductive number and generation interval. To illustrate these approaches, we present them in the context of two influenza A virus (IAV) ferret transmission experiments: one using influenza A/California/07/2009 (Cal/2009) and one using influenza A/Hong Kong/1/1968 (Hong Kong/1968). Despite estimating broadly similar transmissibilities for Cal/2009 and Hong Kong/1968, we conclude that Cal/2009 has a higher pandemic potential. This difference in pandemic potential seems to be primarily driven by its more robust within-host replication. Our results critically depend on several assumptions, namely that the within-host viral dynamics in humans and those in the model animal used (here, ferrets) share quantitative similarities and that viral transmissibility between model animals reflects viral transmissibility between humans. The methods we present to assess relative pandemic risk across viral isolates can be used to improve quantitative risk assessment of other emerging viruses of pandemic concern.

**Author summary:** Pandemic viruses often originate in animal reservoirs. Surveillance studies at the animal-human interface try to identify which of the many viruses circulating in animals could cause a pandemic in humans. Experimental transmission studies in animal models are one such method used to characterize potential pandemic viruses. However, only simple outcomes usually get reported from these experiments, such as the fraction of animals that become infected in such an experiment. Here, we develop a framework to extract more information from these experiments. Specifically, we use data on the virus population within infected ferrets to extrapolate information about transmission. We then simulate many transmission events to estimate epidemiological parameters commonly used in public health to quantify pandemic potential. Our methods therefore allow us to use commonly gathered data from experimental transmission studies to estimate epidemic dynamics in humans. This set of methods can be used to compare the relative pandemic risk of animal viruses before they emerge in humans.

## Introduction

Pandemics arising via zoonotic spillover from animal reservoirs have resulted in mass human mortality and extreme societal disruption. Notable examples are the 1918 Spanish flu pandemic caused by influenza A virus (IAV) H1N1 and the COVID-19 pandemic caused by SARS-CoV-2. Given the impact of pandemics, there is considerable interest in preventing pathogen emergence, particularly of rapidly spreading respiratory pathogens. One key component in preventing pathogen emergence involves surveillance of reservoir species to identify viruses with pandemic potential. However, reservoir species can harbor diverse viral pathogens, resulting in a large pool of possible pre-pandemic strains. Approaches to winnow down the considerable diversity of candidate viruses to those that are more likely to cause a pandemic are therefore essential to guiding control efforts to mitigate pandemic risk. One approach for prioritizing viruses for preparedness and control strategies involves genotypic and phenotypic characterization of viral strains to identify those with traits associated with pandemic risk [1, 2]. For example, assays measuring IAV hemagglutinin receptor binding specificity are used to determine if an influenza virus preferentially binds cells with *α*-2,3 or *α*-2,6 sialic acid linkages. IAVs with the *α*-2,3 binding preference generally do not transmit well from person to person. For many relevant traits, however, genotypic characterizations provide only limited information because the relationships between viral genotypes and their phenotypes have not been fully characterized. Risk assessments based on genetic analyses are therefore limited to traits where the genetic basis is well understood.

Experimental transmission studies have also been widely used to estimate the transmission potential of viral strains. For IAV risk assessment, ferrets are often used because of similarities between ferrets and humans in phenotypes relevant to infection and transmission, such as viral attachment [3] and clinical signs. A notable study found evidence for a strong positive correlation between ferret-to-ferret transmission probability (as measured by secondary attack rate) and IAV circulation patterns in humans [4]. The findings of this study were particularly impactful because numerous ferret experiments were used in the assessment of the relationship between secondary attack rates and IAV circulation patterns in humans. Results from these types of transmission experiments provide semi-quantitative measurements of transmission potential, one aspect of pandemic risk. However, the read-outs of these experiments are typically low-resolution estimates. For example, in a study with three index-contact pairs, the observed overall transmission efficiency could be 0% (0/3), 33% (1/3), 67% (2/3), or 100% (3/3).

Given the cost and effort involved in carrying out these experiments, extracting more information about transmission from these data would be worthwhile. To this end, we here develop new methods to improve risk assessment using measurements commonly taken during experimental transmission studies. We specifically propose to use within-host viral titer data to improve estimation of transmission potential and pandemic risk assessment. Our approach relies on the titer data from infected index animals as well as infected contact animals to provide quantitative estimates of transmission parameters and epidemiological components of pandemic risk. To motivate and illustrate our approach, we apply it to data from two recently performed IAV experimental transmission studies [5], the first using a 2009 pandemic H1N1 isolate and the second using a 1968 pandemic H3N2 isolate. Below, we first present the data from the experimental transmission studies and then develop our quantitative approaches in application to these data.

## Results

### Observed infection dynamics in the experimental IAV transmission studies

The airborne transmission studies analyzed here have previously been described in detail, and their data have been made publicly available [5]. A brief overview of the study design is provided in the Methods section. In these studies, airborne transmission was assessed from index ferrets to contacts using inoculation doses from 10^0^ to 10^6^ TCID_50_. We considered an animal to be infected if viral shedding over the limit of detection (10^1^ TCID_50_/mL) was observed at any sampled timepoint. Transmission efficiency was defined as the fraction of contact ferrets that were infected by their paired index in the subset of pairs in which the index became infected.

The transmission efficiency for each experiment is listed in Figure 1. For influenza A/California/07/2009, it ranges from 75-100%; for influenza A/Hong Kong/1/1968, it ranges from 33-100%. Interestingly, the transmission efficiencies observed for Cal/2009 all fall within the range of respiratory droplet attack rates that Buhnerkempe and colleagues [4] would predict as having a high probability of a supercritical classification in humans. In contrast, the efficiencies observed in the Hong Kong/1968 transmission study span a range between subcritical and supercritical classification in humans.

**Figure 1.**
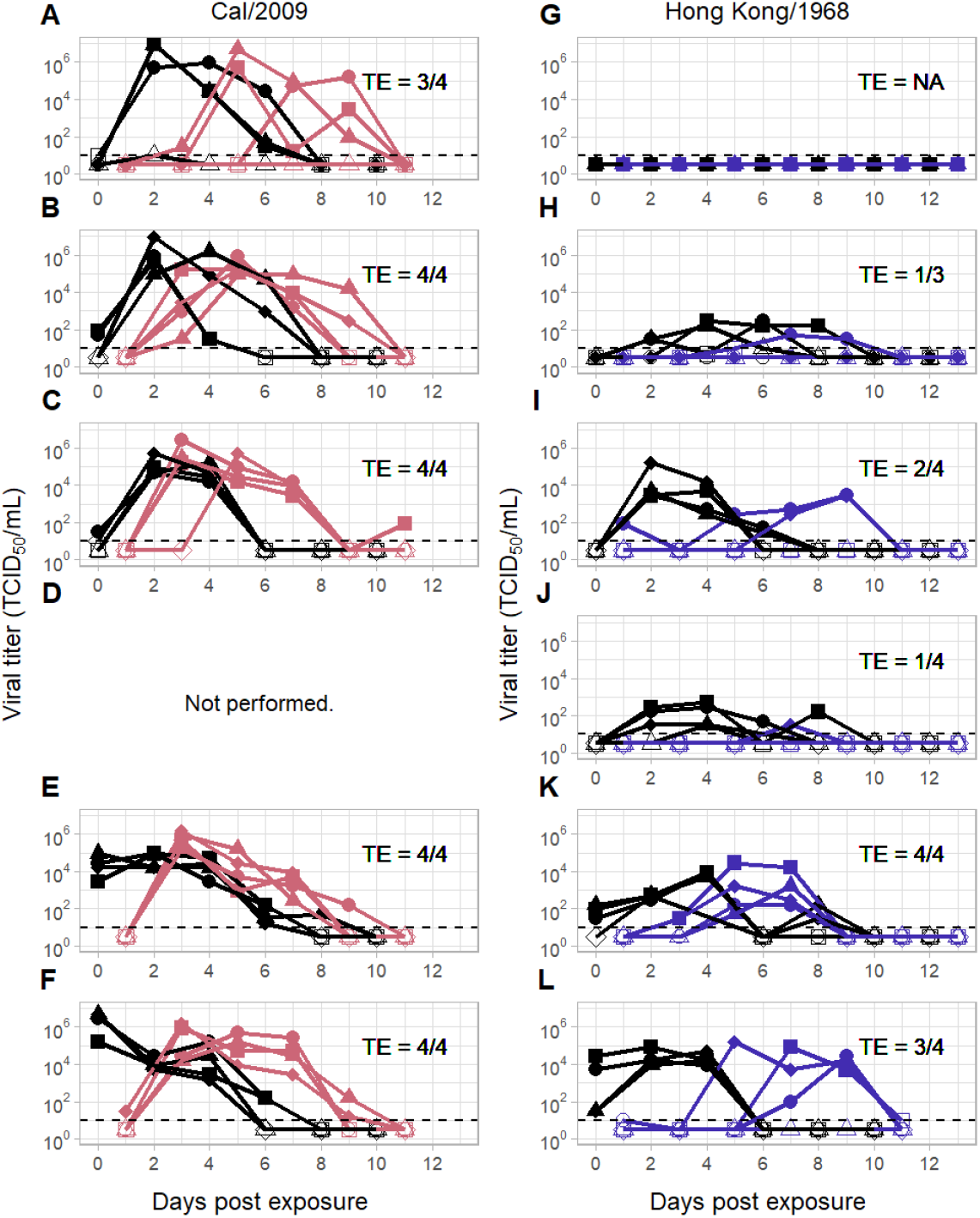
Viral kinetics in the Cal/2009 and Hong Kong/1968 experimental transmission studies. The left column shows Cal/2009 viral kinetics in index ferrets (black) and respiratory contact ferrets (pink). The right column shows Hong Kong/1968 viral kinetics in index ferrets (black) and respiratory contact ferrets (blue). Rows correspond to different index inoculum doses: (A, G) 10^0^, (B, H) 10^1^, (C, I) 10^2^, (D, J) 10^3^, (E, K) 10^4^, and (F, L) 10^6^ TCID_50_*/*mL. The limit of detection is shown as a dotted line at 10^1^ TCID_50_/mL. Transmission pairs share the same marker. Open markers at 10^0.5^ TCID_50_/mL denote samples with viral titers that fall below the limit of detection. Each panel further provides the transmission efficiency (TE) in that experiment, with the denominator denoting the number of infected index animals in the experiment and the numerator denoting the number of contact animals that became infected. NA = Not Applicable. Data are from [5].

### Estimation of the transmission function

Several qualitative patterns emerge from inspection of the viral titer dynamics shown in Figure 1. First, the probability of transmission to a contact animal generally appears to be lower when viral titers in the index are lower. Second, particularly in the Cal/2009 study, transmission to a contact animal appears to occur earlier at higher inoculum doses. This is likely because higher inoculum doses tend to result in higher viral titers in the index animals shortly following challenge. As such, viral titer levels appear to impact not only whether a contact animal gets infected but, in the case of infection, when it gets infected. This observation is key to the development of our analytical approach.

Given that the viral kinetics of the index appear to impact the probability and timing of transmission, we here develop a quantitative framework for estimating viral transmissibility between index and contact animals while accounting for viral titers. We specifically rely on approaches that allow for a time-varying force of infection to quantify how the transmission rate between an index animal and its contact depends on time-varying index viral titers.

We define the force of infection *λ* that an index animal exerts on a contact animal at time *t* as some functional form that depends on the index animal’s viral titer at time *t*. For example, we can let *λ* linearly depend on the *log*_10_ of the index animal’s viral titer based on empirical support for this functional relationship from previous studies [6, 7, 8]. In this case, the force of infection is given by:

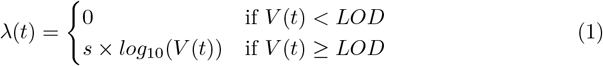

where viral load measured at time *t* (*V* (*t*)) is measured in TCID_50_/mL, parameter *s* quantifies transmissibility, and LOD denotes the limit of detection of the assay used to quantify viral titers. The probability that an index transmits to its contact and the timing of the transmission event are both functions of this time-varying force of infection (Methods). Given observed index viral titers (Figure 1) and information on infection times of the contacts, we can statistically estimate the parameter *s* (Methods). Figure 2A shows the estimates of *s* derived from the Cal/2009 and Hong Kong/1968 transmission studies. For Cal/2009, the maximum likelihood estimate is *s* = 0.111 (95% confidence interval (CI) = [0.068, 0.172]) and for Hong Kong/1968, it is *s* = 0.047 ([0.024, 0.082]). Our estimates for *s* indicate that, in infected ferrets, the transmissibility of Cal/2009 tends to be higher than that of Hong Kong/1968 while controlling for index viral titers. To determine whether the transmissibility *s* of Cal/2009 is statistically higher than that of Hong Kong/1968, we estimated *s* in a model which forced Cal/2009 and Hong Kong/1968 to have the same transmissibility and then calculated the level of support for this simpler model relative to the model with two separate transmissibility values (one for Cal/2009 and one for Hong Kong/1968) (Methods). While we found that the model with two separate transmissibilities was statistically preferred, even when penalizing for model complexity, the extent to which it was preferred was small (Δ*AICc* = 2.74), indicating that the statistical support for the higher transmissibility of Cal/2009 relative to Hong Kong/1968 is not strong.

**Figure 2.**
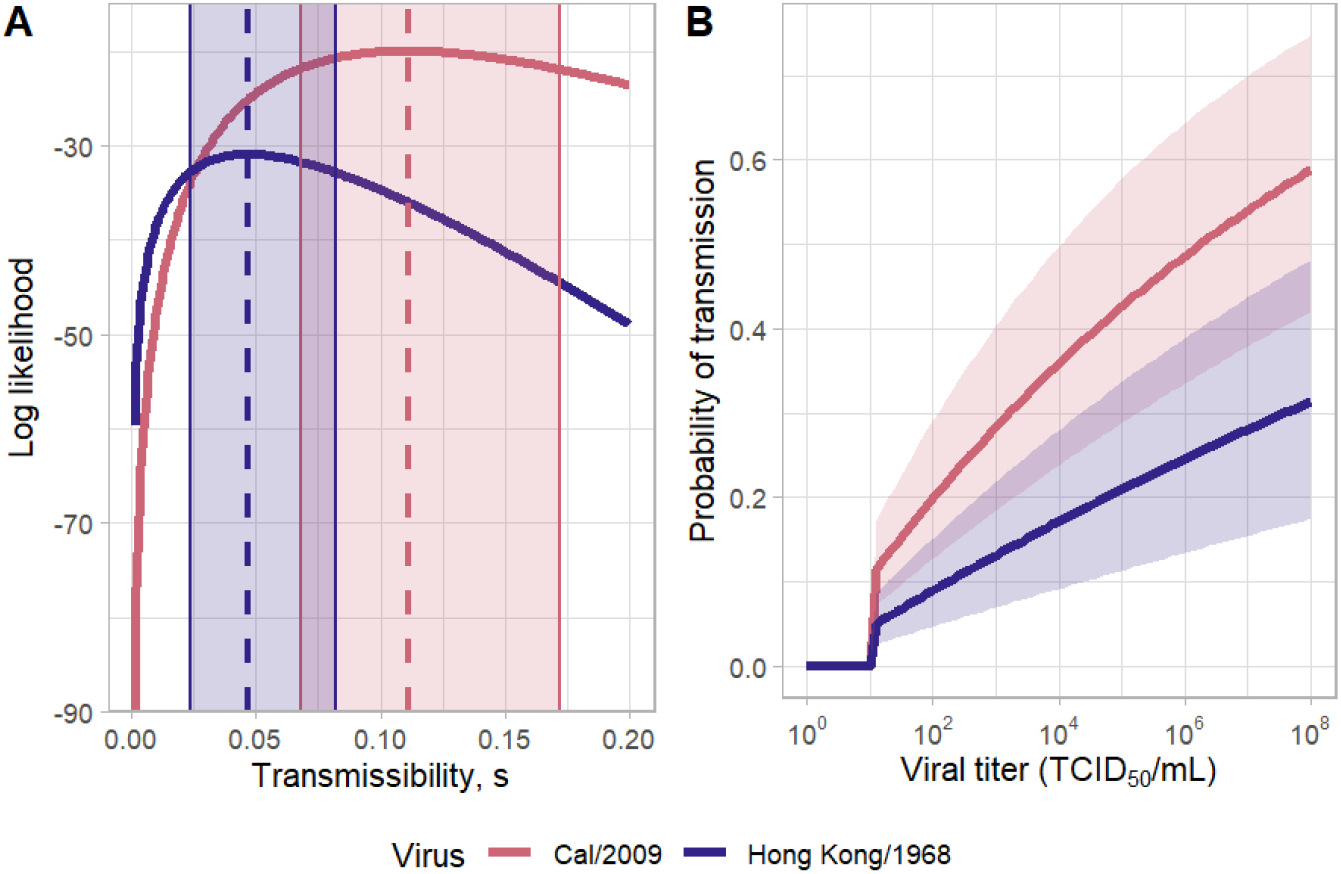
Statistical estimation of the *log*_10_ transmission function. (A) The log likelihood profile for the transmissibility parameter *s* for both the Cal/2009 virus (pink) and the Hong Kong/1968 virus (blue). Dashed lines show the maximum likelihood values of *s*. Shaded regions show the 95% confidence intervals. (B) Calculated probabilities of transmission to a contact after a one-day exposure to an index animal with a constant viral titer, as given on the x-axis. Solid lines show the probabilities calculated using the maximum likelihood estimates for *s*. Shaded regions show transmission probabilities that fall within the 95% confidence interval values for *s*.

To gain better intuition into the extent to which the inferred transmissibilities of Cal/2009 and Hong Kong/1968 differed from one another, we calculated the probability of a contact becoming infected if exposed for a period of one day to an index animal with a given viral titer (Figure 2B). Given the values of *s* for Cal/2009 than for Hong Kong/1968, the probability that a contact becomes infected during this day of exposure is predicted to be higher (approximately twice as high) for Cal/2009 than for Hong Kong/1968 at a given index viral titer.

### Assessment of alternative transmission function forms

Because we do not a priori know that the *log*_10_ functional form for the force of infection that we used in equation (1) is an appropriate one to use, we next considered alternative functional forms for the relationship between force of infection *λ* and viral titers. The alternative functional forms we considered were a basic linear relationship, a threshold relationship (whereby *λ* was zero below a threshold viral titer and some constant value above that threshold titer), and a flexible Hill function (Methods). We estimated the parameters for each of these models for each viral isolate. Figure 3 graphically shows these force of infection estimates for the *log*_10_ model and the alternative models for each of the two viral isolates, along with the calculated probabilities of transmission given a one-day exposure period to an index animal with a given viral titer. To determine which of these models performed best, we then calculated the AICc scores for each model (Table 1). The linear form performed very poorly for both Cal/2009 and Hong Kong/1968. The threshold model had the best AICc score for both isolates, but there is little statistical support for discriminating between the *log*_10_, threshold, and Hill forms. Comparing the force of infection and probability of transmission results in Figure 3 for these three forms demonstrates that the results (at maximum likelihood parameterization) are quite similar, particularly between the threshold and Hill functional forms.

**Table 1.** Model comparison of original *log*_10_ and alternative functional forms for the force of infection. The columns specify the model name, the model parameters, the corrected AIC (AICc) value for the model fit to the Cal/2009 viral isolate data, and the corrected AIC value for the model fit to the Hong Kong/1968 viral isolate. Models with lower AICc are preferred. ΔAICc values are also shown in parentheses for both viral isolates, relative to the statistically most preferred model, whose AICc values are shown in boldface.

|  | Parameters | Cal/2009 AICc | Hong Kong/1968 AICc |
| --- | --- | --- | --- |
| Linear | $s_L$ | 154.5 (116.05) | 122.59 (59.46) |
| Log-transformed | $s$ | 42.22 (3.77) | 64.12 (0.99) |
| Threshold | $h, r$ | **38.45** | **63.13** |
| Hill | $q, k_a, n$ | 42.47 (4.02) | 66.05 (2.92) |

**Figure 3.**
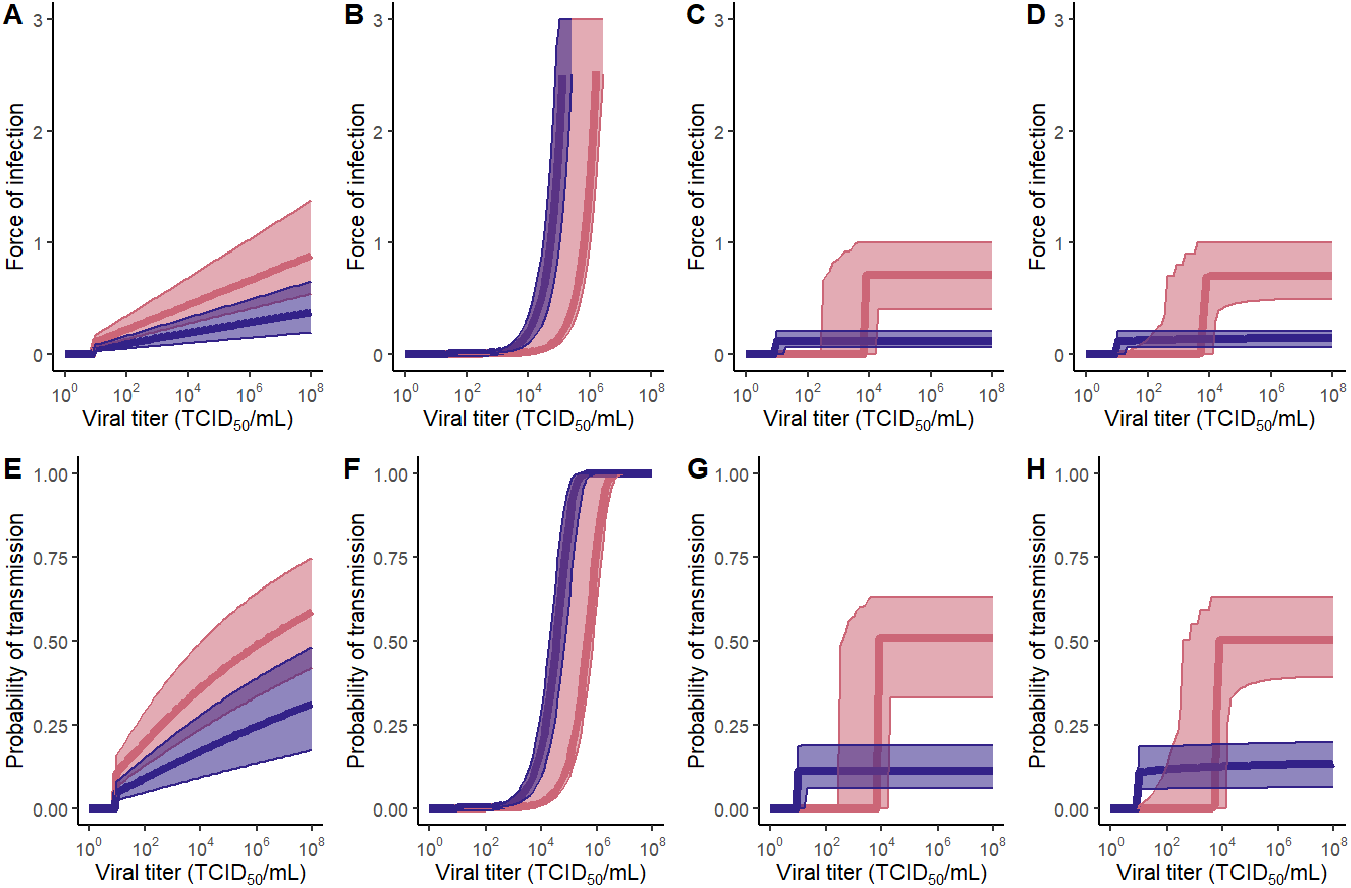
Transmissibility estimates for alternative force-of-infection functional forms. (A)-(D) Estimated forces of infection (*λ* values) across a range of viral titers. (E)-(H) Probabilities of transmission given a one-day exposure to a constant viral titer, given the *λ* estimates from (A)-(D), respectively. Columns correspond to alternative functional forms: (A,E) *log*_10_-transformed, (B,F) linear, (C,G) threshold, and (D,H) Hill function. The maximum likelihood estimates for Cal/2009 are *s*_*L*_ = 1.60*×* 10^−6^ for the linear function (B,F), *r* = 0.71 and *h* = 3.83 for the threshold function (C,G), and *q* = 0.7 *k*_*a*_ = 3.8, and *n* = 198.8 for the Hill function (D,H). The maximum likelihood estimates for Hong Kong/1968 are *s*_*L*_ =1.99 *×* 10^−5^ for the linear function (B,F), *r* = 0.12 and *h* = 1.0 for the threshold function (C,G), and *q* = 0.3, *k*_*a*_ = 6000, and *n* = 0.012 for the Hill function (D,H). Shaded regions show 95% confidence intervals.

### Quantification of parameters at the transmission event

Whereas the index animals in the Cal/2009 and Hong Kong/1968 experimental transmission studies received direct intranasal challenge with varying inoculum doses, the contact animals acquired their infections via inhalation of infectious respiratory particles that were shed by the index animals. As such, one could argue that the viral titer dynamics observed in the contact animals are more likely to resemble natural infections. Indeed, a qualitative re-inspection of Figure 1 indicates that, for Cal/2009, neither the peak viral titer nor the duration of infection in contact animals appears to depend on the index’s inoculum dose. Similarly, in the Hong Kong/1968 study, although there appears to be considerably more heterogeneity across the contact animals in their viral titer dynamics, neither peak viral titers nor infection durations appear to depend systematically on index inoculum doses or their viral kinetics. In both studies, viral titers in the contact animals appear to start off at low levels, reaching their peaks within 2 to 4 days. Here, to quantify parameters at the transmission event, we therefore assume that the viral titer dynamics observed in the contact animals are representative of those in natural infections. Moreover, for projecting relative transmission potential, we need to assume that the contact animal viral dynamics are not only similar to those of natural ferret infections with the isolates considered, but also similar to those of natural human infections with the isolates considered. This is not a trivial assumption, and we return to this point in the Discussion.

Our approach to projecting parameters across the transmission event relies on combining viral titer dynamics observed across contact ferrets with our parameterized transmission function. These contact ferrets become “theoretical” index animals, and we use their observed titer dynamics and the transmission function to simulate the expected number of secondary infections (Methods). Figures 4A and 4B show a single stochastic realization of the offspring distribution for Cal/2009 and Hong Kong/1968 “theoretical” index animals, respectively, assuming an average of 15 hour-long contacts per day and using the *log*_10_ transmission function, parameterized with *s* values randomly drawn from the estimated 95% confidence interval of this parameter. We chose a rate of 15 contacts per day based on contact rate estimates for children under the age of 18 [9]. We chose the *log*_10_ transmission function due to its support from previous studies and its good performance relative to the alternative models considered (Table 1). By eye, both offspring distributions appear to follow a negative binomial distribution with different means. From our stochastic simulations, we could also keep track of when transmissions occurred from each infected animal and use this information to calculate the generation interval, defined as the time between the infected animal’s infection and onward transmission. Figures 4C and 4D show these generation times for the Cal/2009 and Hong Kong/1968 secondary infections that were realized and shown in 4A and 4B, respectively.

**Figure 4.**
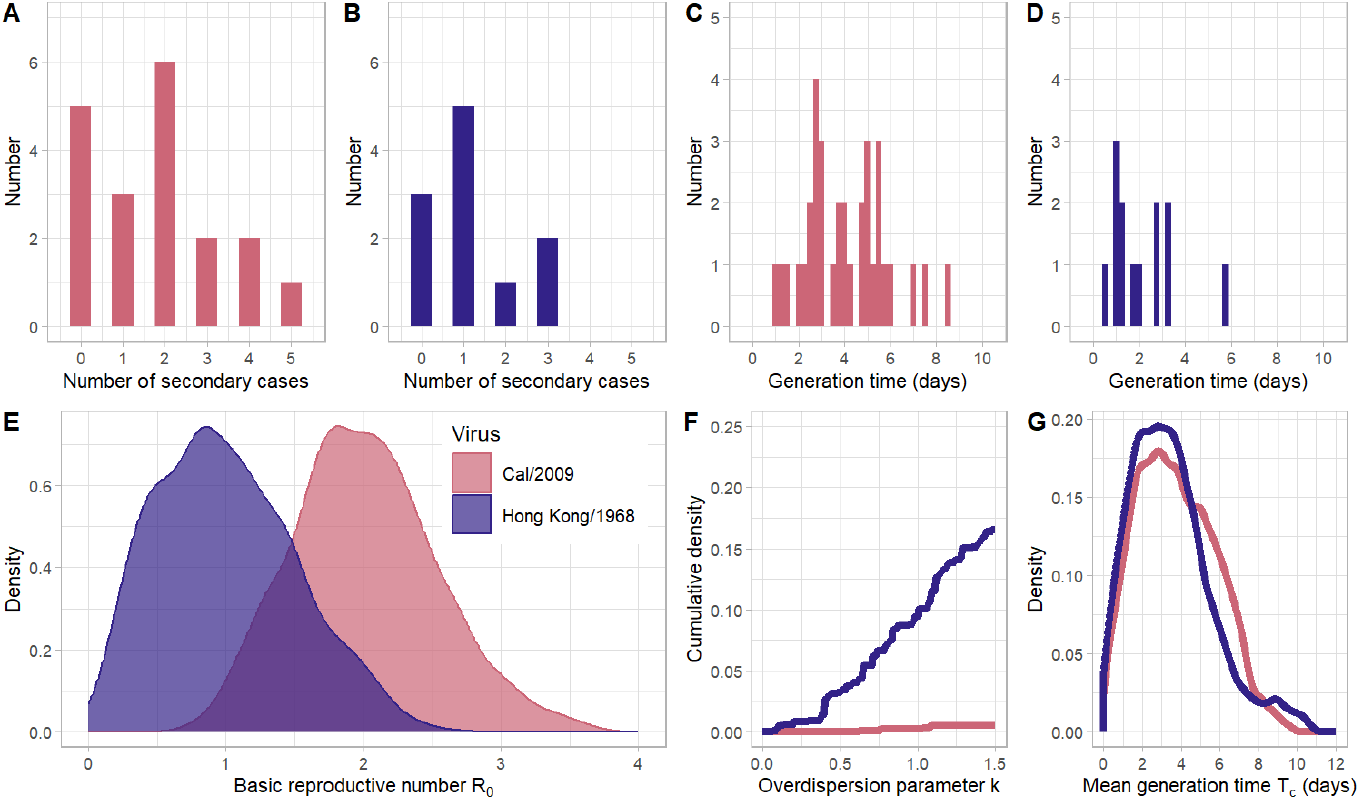
Estimation of transmission event parameters for Cal/2009 and Hong Kong/1968. (A) Distribution of the number of secondary infections generated by *n*=19 Cal/2009 contact animals. (B) Distribution of the number of secondary infections generated by *n*=11 Hong Kong/1968 contact animals. In panels (A) and (B), the distributions each show the outcome of one stochastic realization, assuming each animal has 15 one-hour long contacts per day. (C) The generation intervals for each secondary case (*n*=34) generated in Cal/2009 simulation shown in panel (A). (D) The generation intervals for each secondary case (*n*=13) generated in the Hong Kong/1968 simulation shown in panel (B). (E) The distribution of *R*_0_ values from the 1000 Cal/2009 and 1000 Hong Kong/1968 stochastic simulations. (F) The cumulative distribution of overdispersion (*k*) values from the Cal/2009 and Hong Kong/1968 stochastic simulations that resulted in at least one individual transmitting infection. This was 1000 simulations for Cal/2009 and 997 simulations for Hong Kong/1968. (G) The distribution of mean generation intervals from the 1000 Cal/2009 and Hong Kong/1968 stochastic simulations.

To scale up from the single stochastic realizations shown in 4A-D for the populations of Cal/2009 and Hong Kong/1968 “theoretical” index ferrets, we performed 1000 stochastic realizations for each virus. To ensure that our simulations incorporated uncertainty in the value of *s*, we sampled values of *s* from the likelihood profile we identified previously (Figure 2A) using a Metropolis-Hastings algorithm (Methods). Each of the 1000 stochastic simulations used a *s* parameter drawn with a probability equal to its likelihood (Methods). Each of the 1000 simulations included all of the contact animals that were infected in the original study (*n*=19 for Cal/2009 and *n*=11 for Hong Kong/1968). For each realization, we fit a negative binomial distribution to the resultant offspring distribution and calculated the mean and the overdispersion parameter *k* of this distribution. The mean of this distribution is, by definition, equivalent to the reproduction number *R*_0_, estimated from a single stochastic realization. Figure 4E plots the distribution of *R*_0_ values that were estimated for the Cal/2009 and Hong Kong/1968 virus isolates using the 1000 stochastic realizations. From this figure, it is clear that the *R*_0_ estimated for the Cal/2009 virus is substantially higher than the one estimated for the Hong Kong/1968 virus, even with considerable uncertainty in its exact value (Welch’s two sample t-test, *p <* 0.001). Moreover, at the contact rate assumed, the *R*_0_ estimated for Cal/2009 exceeds one in almost all of the simulations, whereas it exceeds one for Hong Kong/1968 in fewer than half the simulations. This indicates that, at this contact rate, Cal/2009 would have pandemic potential whereas Hong Kong/1968 would not.

Figure 4F shows the cumulative distributions of the overdispersion parameters estimated for Cal/2009 and Hong Kong/1968 from their respective stochastic realizations. For Cal/2009, we find that 2 of the 1000 stochastic simulations have an estimated overdispersion parameter *k* of less than one. For Hong Kong/1968, around 10% of simulation have an estimated *k* of less than one. This indicates that there is relatively little transmission heterogeneity for both viral isolates, with transmission heterogeneity being particularly low for Cal/2009. Note that, for both viral isolates, the estimated overdispersion values only capture transmission heterogeneity stemming from interindividual variation in viral titers. As such, our estimates of the extent of transmission heterogeneity will be lower-bound estimates. This is because of additional variation that contributes to transmission heterogeneity, such as heterogeneity in contact rates that occur in real world settings.

Finally, we calculated the mean generation interval *T*_*c*_ for Cal/2009 and for Hong Kong/1968 from each of the stochastic simulations. The distributions of these mean generation intervals are shown in Figure 4G. We find that the mean generation interval for Hong Kong/1968 is slightly shorter than for Cal/2009, at 3.47 verses 3.78 days (Welch’s two sample t-test, *p <* 0.001). However, this difference is unlikely to be biologically significant.

In sum, for both of the viral isolates considered, we calculated three epidemiological parameters that are relevant to pandemic risk: the basic reproduction number (under an assumption of a contact rate of 15 individuals per day), the overdispersion parameter, and the generation interval. Generally, we expect that pandemic risk is greater for virus isolates with higher *R*_0_, low overdispersion, and short generation intervals (due to more rapid spread). As such, a head-to-head comparison of Cal/2009 against Hong Kong/1968 would indicate that Cal/2009 would pose greater pandemic risk than Hong Kong/1968.

### Prediction of pandemic potential and dynamics

Above, we assumed a single contact rate (an average of 15 hour-long contacts per day) to project the number of secondary infections for each of the two viruses and to estimate their generation intervals. The chosen contact rate impacts the distribution of secondary infections and thus our estimates of *R*_0_. In Figure 5A, we show how our *R*_0_ estimates change for Cal/2009 and Hong Kong/1968 across a broad range of contact rates. The range of contact rates we consider span those deemed relevant for a respiratory virus of humans [9].

**Figure 5.**
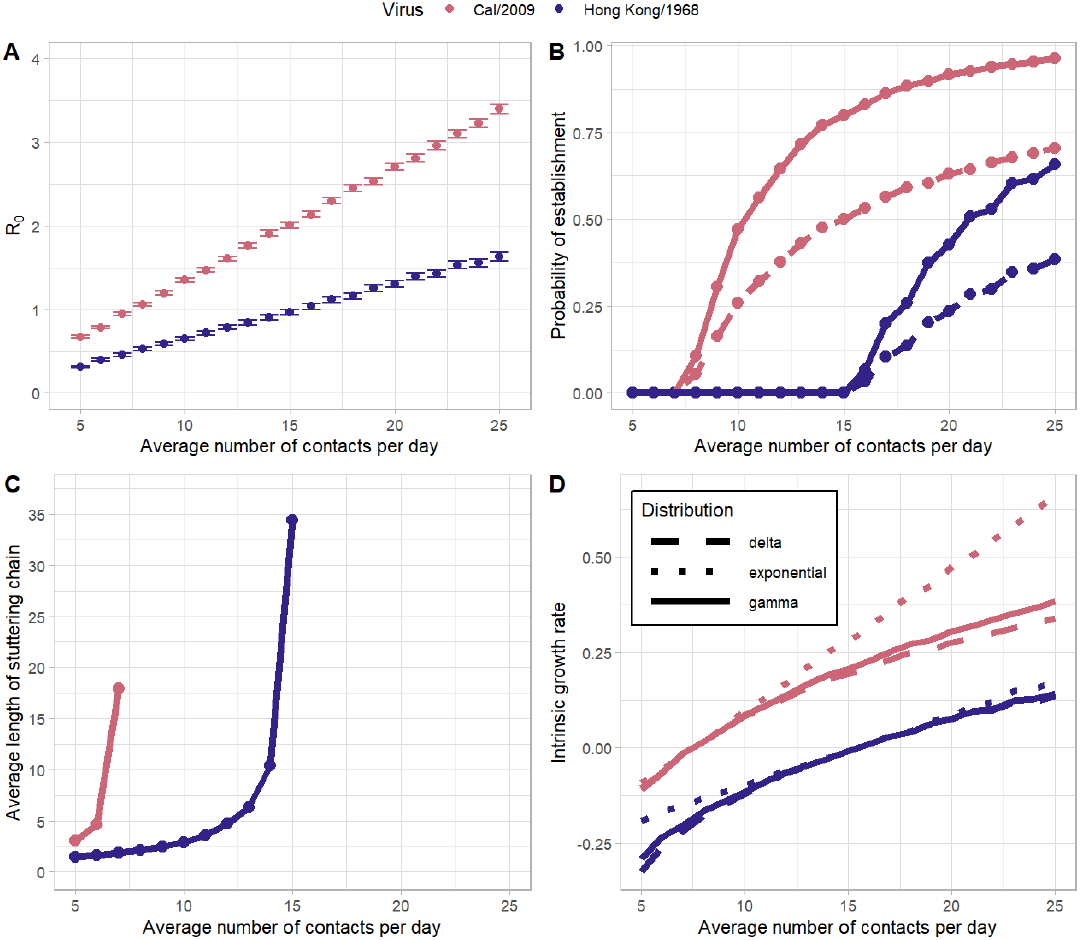
Prediction of pandemic potential and dynamics. (A) Estimates of the reproduction number *R*_0_ for Cal/2009 and Hong Kong/1968 across a range of contact rates. Error bars indicate the 95% confidence intervals. (B) The probability of pandemic establishment for Cal/2009 and Hong Kong/1968 across a range of contact rates. The solid lines indicate probabilities in the absence of transmission heterogeneity (*k* = *∞*). The dashed lines indicate probabilities with a moderate level of transmission heterogeneity (*k* = 1). (C) The average length of stuttering transmission chains across the range of contact rates when *R*_0_ *<* 1 for both viruses. (D) Estimated intrinsic growth rates (per day) of Cal/2009 and Hong Kong/1968 across a range of contact rates, assuming an exponential (dotted lines), delta (dashed lines), or gamma (solid lines, *n* = 5) distribution of generation intervals. The doubling time of an epidemic decreases as *r* increases. When *r* = 0.07, the doubling time is 10 days; when *r* = 0.14, the doubling time is 5 days; and when *r* = 0.35, the doubling time is 2 days. In all panels, Cal/2009 projections are shown in pink and Hong Kong/1968 projections are shown in blue.

We can use the *R*_0_ estimates shown in Figure 5A to predict the risk of pandemic establishment of the two viruses, given an assumed contact rate (Methods). Figure 5B shows this establishment risk for each virus, under two different transmission heterogeneity assumptions: one with no transmission heterogeneity (*k* = *∞*) and one with a moderate level of transmission heterogeneity (*k* = 1) that exceeds our simulated expectations (Figure 3F). For Cal/2009, at contact rates below 8 hour-long contacts per day, the virus is not predicted to establish due to a subcritical basic reproduction number (*R*_0_ *<* 1). For Hong Kong/1968, the first contact rate at which *R*_0_ exceeds 1 is 16 contacts per day. Increases in daily contact rates above these critical values result in higher probabilities of viral establishment, although the exact probabilities depend on the extent of transmission heterogeneity. As theoretically expected [10], higher levels of transmission heterogeneity lower the probability of viral establishment. In the absence of transmission heterogeneity (*k* =*∞*), the probability that Cal/2009 establishes is approximately 89% when the contact rate is 18 contacts per day, corresponding to the contact rate of the age group with the highest number of daily contacts in the study by Mossong and colleagues [9]. At a contact rate of 7 contacts per day, corresponding to the contact rate of the age group with the lowest number of daily contacts, the probability that Cal/2009 establishes is 0%. Likewise, for Hong Kong/1968, the probability of establishment is 26% at 18 contacts per day and 0% at 7 contacts per day.

Previous studies have pointed out that initial *R*_0_ values for a spillover virus matter for projecting pandemic risk, even if they fall below one [11]. This is because viruses have the potential to adapt as they transmit between individuals along stuttering chains. At *R*_0_ values closer to one, the length of these stuttering chains is expected to be longer than at *R*_0_ values closer to zero, providing viruses with higher *R*_0_ greater opportunity to adapt and thus to ultimately establish. In Figure 5C, we plot for each virus considered, the average length of stuttering chains across the assumed range of contact rates (Methods). We plot these average lengths only for the subset of contact rates for which a virus still has a subcritical (*<* 1) *R*_0_. At higher contact rates, the average length of stuttering chains is higher.

Finally, we can use our estimates of *R*_0_, shown in Figure 4A, along with a specified generation time distribution, to estimate the intrinsic growth rates (*r* values) of the viruses under various contact rate scenarios. The intrinsic growth rate of a virus is important to project in that it specifies the viral doubling time at the level of the host population and therefore the speed at which a pandemic takes off [12]. Figure 5D shows intrinsic growth rate estimates for both viruses across the range of assumed contact rates, under the assumption of a mean generation time of 3.6 days (Methods). Because estimates of the intrinsic growth rate are impacted by the shape of the generation interval distribution (not only its mean) [13, 14], we consider three parameterizations of a gamma distribution for the generation interval distribution: one corresponding to an exponential distribution (where the variance of the distribution is equal to the mean of the distribution), one corresponding to a delta distribution (where all probability density lies at the mean value), and one that is empirically more realistic, lying between these two extremes. For both viruses, projected intrinsic growth rates are higher at higher contact rates, as expected. At contact rates that yield supercritical *R*_0_ values, intrinsic growth rates are higher (and doubling times shorter) under the exponential distribution than under the delta distribution, again, as expected. The empirically realistic gamma distribution predicts intrinsic growth rates that fall between these two extremes, closer to the rates predicted by the delta distribution.

In sum, for both of the viral isolates considered, we quantified several metrics relevant to pandemic risk across a range of contact rates: the probability of epidemic establishment, the average length of stuttering transmission chains, and the intrinsic growth rate of an epidemic. Generally, we expect that virus isolates with *R*_0_ values exceeding one across a broad range of reasonable contact rates, those with high probabilities of establishment, those with long stuttering transmission chains, and those with higher intrinsic growth rates all have an elevated pandemic risk. A head-to-head comparison of Cal/2009 against Hong Kong/1968 indicates that Cal/2009 would pose greater pandemic risk than Hong Kong/1968.

## Discussion

Here, we developed an analytical approach to gauge the pandemic potential of zoonotic viruses using viral titer data from experimental transmission studies. Our approach relied on estimating the parameters of a transmission function using information on index animal viral kinetics and the timing of contact animal infection. It further relied on assuming that the viral kinetics of contact animals reflected those of animals that would be naturally infected with the viruses considered, and more stringently, those of humans who would become naturally infected with the zoonotic viruses considered. We applied our approach to existing transmission study datasets that used two IAV isolates and found that Cal/2009 had higher pandemic potential than Hong Kong/1968 (Figures 4 and 5). We imagine the primary use case for this approach would be in assessing the relative pandemic potential of animal-origin influenza viruses which have not yet emerged in humans. These results, and indeed the results of any subsequent analyses using this method, are valid only for the specific isolates used in the experimental studies. The continual and rapid evolution of influenza viruses precludes us from generalizing these results to other H1N1 or H3N2 strains, including those which will arise in the future and could potentially cause pandemics.

We now turn to discussing some of the benefits that come with the analytical approach we presented here. In addition to being able to assess relative pandemic potential of different viral strains, our approach provides insight into why one virus might have higher transmission potential than another. For example, we found that our estimate of the transmissibility parameter *s* was slightly higher for Cal/2009 than it was for Hong Kong/1968 (Figure 2). This indicates that an individual infected with Cal/2009 at a given viral titer is more likely to transmit to a contact than an individual infected with Hong Kong/1968 at the same viral titer. However, it seems likely that the primary reason that Cal/2009 had a higher transmission potential than Hong Kong/1968 was because viral titers were higher in Cal/2009-infected animals than in Hong Kong/1968-infected animals (Figure 1). Mutations which increased viral titers in Hong Kong/1968 infected individuals to the level of Cal/2009 would close the gap in transmission potential between these viruses. In contrast, if the primary reason for Cal/2009 having higher transmission potential than Hong Kong/1968 was due to higher transmissibility (higher *s*), then one might expect that mutations that result in higher transmissibility might be selected for in zoonotic Hong Kong/1968 viral populations. Our approach therefore has the potential to help disentangle factors contributing to pandemic potential. An interesting application of this ability would be to consider viral strains differing by a handful of mutations, for example an early pandemic strain and one that carries adaptive mutations from circulation in humans for some relatively short period of time. With data from these types of experimental transmission studies, one could identify whether the observed mutations increased transmissibility (that is, increased *s*) or simply increased viral titers that would impact downstream transmission. We next used *s* to estimate *R*_0_, an epidemiological parameter commonly used to assess pandemic risk. Because there are many factors relevant to transmission which are not captured by the experimental setup, our estimates of *R*_0_ should be interpreted as relative values. As such, we cannot interpret these values as being sub- or supercritical (that is, *R*_0_ is less than or greater than 1). We conduct all subsequent analyses (Figure 5) across a range of contact rates to account for this uncertainty.

We aim to use experimental transmission studies in animal models to assess relative pandemic risk in humans. There are several intrinsic limitations associated with this goal, independent of the specifics of the analytical approach. Most obviously, ferrets are not humans. Results based on transmission experiments in ferrets are only relevant to assessing pandemic risk for humans if transmission potential in ferrets reflects transmission potential in humans. One way to circumvent this is to analyze experimental transmission studies in humans [15]. However, the number of transmissions will be severely limited in the case of human studies, such that it is unlikely that transmission parameters could be estimated with confidence. Additionally, it is unethical to test transmission of emerging viruses in humans because of possible adverse events and side effects. As such, experimental transmission studies in model animals such as ferrets remain the most promising for assessing pandemic risk in humans. Despite the limitations associated with ferrets as a model system, previous analyses do indicate that human infection risk is positively associated with secondary attack rates in ferrets [4], mitigating some of the concern that transmission potential in ferrets does not correlate with transmission potential in humans. Furthermore, subject matter experts generally agree that animal transmission experiments are important for gauging pandemic risk in humans, which is why this metric has been incorporated into frameworks such as the CDC Influenza Risk Assessment Tool [16].

Experimental transmission studies are additionally limited by sampling limitations. In transmission studies like those analyzed here, nasal washes or swabs are often taken from a ferret every or every other day. Viral titers between these measured time points are then inferred through interpolation between the available data points. The viral titers we use to estimate transmissibility therefore do not fully capture the entirety of viral population dynamics in infected hosts. For example, it is unlikely that the time of peak viral titer in an animal coincides with the time that a sample is taken from that animal. Our method is therefore likely to systematically overestimate transmissibility (s), as our interpolations miss the viral peaks. Sampling intervals and assay limits of detection also affect when we can say that transmission from index to contact has occurred. If sampling was performed more frequently or assay limits of detection were lower, times of transmission would be more finely resolved. Moreover, our inference approach currently assumes that the contact is still uninfected on the last negative test day (*T*_1_) prior to the first positive test day (*T*_2_). However, this does not necessarily need to be the case: a contact could already be infected on *T*_1_ but with viral titers that fall below the limit of detection. Lower limits of detection would thus improve our inference. Improvements in inference could also come from computational extensions of our approach that would allow for the possibility of infection prior to detection. Our model can be further extended to consider measurement noise in viral titer data and stochastic viral dynamics early on during infection.

A second limitation is that our approach assumes that transmission potential is related to viral titer levels in an index, as measured by nasal wash. If transmission potential depended more strongly on viral titers in other locations, our results of relative pandemic risk could be biased. However, several studies using genetically modified viruses have shown that H1N1 and H3N2 viruses transmitted through the air predominately originate from the upper respiratory tract, particularly the nasal respiratory epithelium [17]. Thus, we feel confident that the titer measurements we used to estimate the transmission function are taken from biologically relevant locations in the ferret respiratory tract. However, our model in principle could be extended to consider the impact of symptoms on transmissibility, for example, using previously proposed functional forms [18].

Finally, the experimental transmission study we analyzed used ferrets that were naïve to influenza infection. Clearly, cross-immunity generated from previous infection with seasonally circulating viruses could mitigate the risk of pandemic emergence of certain IAV subtypes. To consider the risk of pandemic emergence in the context of existing (cross-)immunity, experimental transmission studies could use ferrets with previous immune histories. Indeed, several transmission studies have already been performed using “pre-immune” ferrets [19] and future work could apply the approach we developed here to data from these studies.

Despite the limitations of our approach and of experimental transmission studies more generally, the method presented here advances integration of pandemic risk assessment with quantitative modeling. We suggest that our approach will be most useful for comparing the pandemic potential of different influenza viruses circulating in non-human reservoirs. Experimental transmission studies are commonly part of an overall risk assessment pipeline that characterizes multiple virological traits thought to be related to pandemic risk [20]. Our data-driven model enriches the insight that can be gained from these experiments. This should help those assessing risk to more accurately weigh pandemic potential and allocate resources appropriately.

Furthermore, we hope that quantitative modeling will become more integrated with efforts to prioritize viruses for enhanced surveillance more generally, whether through experimental challenge and transmission studies or through genetic and phenotypic characterization. A specific benefit of integrating experimental and modeling approaches is the extension of valuable information obtained at the scale of transmission pairs to estimate parameters relevant to viral spread at the population scale. There have been calls for further synthesis of computation and modeling into influenza risk assessments for at least a decade [2], but many aspects of pandemic potential are still studied separately in wet and dry labs. Our method represents one way in which experimental and modeling approaches can be combined for improved influenza pandemic risk assessment.

## Methods

### Design of the experimental transmission study

Here, we analyze data from a series of experimental transmission studies in ferrets carried out in [5]. The first experimental transmission study used the IAV isolate recombinant A/California/07/2009 (Cal/2009). Index ferrets were inoculated at one of five inoculum doses: 10^0^, 10^1^, 10^2^, 10^4^, and 10^6^ TCID_50_. Four index ferrets were challenged at each of these five doses. Each index ferret was housed in a custom-built transmission cage with a single contact ferret. These cages are designed such that the index and contact ferret are separated by a 5 cm wide double offset perforated divider which allows animals to share the same airspace, but prevents direct physical contact. The contact ferret was introduced one day post index inoculation. Nasal washes were collected every other day from index animals beginning on day 1 post inoculation and from contact animals beginning on day 2 post inoculation (1 day post exposure, dpe). For some experiments, samples were collected up to 11 dpe; for others, samples were collected up to 13 dpe. Virus was titrated using tissue culture infectious dose assays with a limit of detection of 10^1^ TCID_50_/mL. For our analyses, the infection status of an index or contact animal was considered positive if at least one sample had a viral titer measurement that fell above the limit of detection (10^1^ TCID_50_/mL).

The second experimental transmission study used the IAV isolate recombinant A/Hong Kong/1/1968 (Hong Kong/1968). Index ferrets were inoculated at one of six inoculum doses: 10^0^, 10^1^, 10^2^, 10^3^, 10^4^, and 10^6^ TCID_50_. Four index ferrets were challenged at each of these six doses. Other components of the Hong Kong/1968 study were analogous to those of the Cal/2009 study.

### Estimation of the transmission function

We quantified the relationship between viral titer and onward transmission potential using longitudinal viral titer data from paired index-contact ferrets. To do this, we use equations developed to estimate the force of infection using the standard ‘catalytic’ model [21]. This model assumes that an individual is initially uninfected and susceptible to infection. With time, the individual may become exposed to the pathogen, such that the probability of testing positive increases with time. More quantitatively, the probability that an individual has been infected by time *T* is given by 1 − e^−*λT*^, where *λ* is a constant force of infection. This equation has been extended to allow for a time-varying force of infection (e.g., see [22]). In this case, the probability that an individual has been infected by time *T* is given by 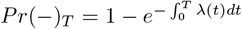 We adopt this time-varying force of infection approach in our analysis of the experimental transmission studies, where the force of infection on the contact animal comes from the index animal. This force of infection is time-varying because it depends on the viral titers of the index animal and these change over time (see Figure 1).

Adopting the *log*_10_ functional form, we let the force of infection be given by equation 1. To evaluate the force of infection *λ*(*t*) from an index ferret over the course of infection, we therefore need to know that animal’s viral titers across all time points. To estimate the complete within-host viral trajectory for each index ferret, we linearly interpolated viral titers on the logarithmic scale. This assumption is equivalent to assuming exponential growth/decay kinetics between measured data points. During this fit, we assume that viral titers below the limit of detection are at half the limit of detection. We further assume that the force of infection *λ* is 0 for all viral titers that fall at or below the limit of detection (LOD) of 10^1^ TCID_50_/mL.

For each contact ferret, we determine whether the ferret has yet been infected with the focal IAV at each of its measured timepoints. Prior to the first positive viral titer measurement, the ferret is considered uninfected. On the day of the first positive viral titer measurement, the ferret is considered infected. We use these timepoints to estimate the parameter *s* of the transmission function using maximum likelihood. For a contact ferret that does not become infected over the study period, we calculate the probability that this contact remained uninfected by the final study timepoint (*T*_*f*_) for a given value of *s*. This probability is given by:

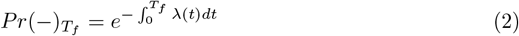

For a contact ferret that does become infected, we compute the probability that this ferret became infected between the time of its last negative test (*T*_1_) and the time of its first positive test (*T*_2_). This probability is given by the product of the probability that the contact has not become infected by time *T*_1_ and the probability that the contact becomes infected between times *T*_1_ and *T*_2_ times:

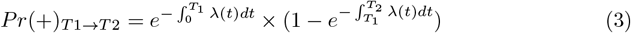

For any value of *s*, the overall likelihood is given by the product of the likelihoods across all contact ferrets (both those who became infected and those that did not). All but one infected index animal was used to estimate *s*. The one index animal (and corresponding contact animal) we excluded was from the Hong Kong/1968 experiment (at a dose of 10^4^ TCID_50_/mL). We excluded this index-contact pair because the index animal was missing one viral titer measurement at 4 days post exposure.

Given a parameterized transmission function, the probabilities of infection shown in Figure 2B are calculated as:

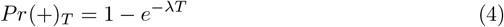

where *T* = 1 day.

To test if transmissibility estimates of *s* were significantly different between Cal/2009 and Hong Kong/1968, we estimated a single value for transmissibility *s* based on both the Cal/2009 and the Hong Kong/1968 data. We then calculated the corrected Akaike information criterion score (AICc) using the *s*_*all*_ maximum likelihood estimate and similarly calculated the AICc using the model with the separate *s* maximum likelihood estimates. The model with separate transmissibilities for Cal/2009 and Hong Kong/1968 had a better (smaller) AICc, however, the Δ*AICc* scores between this model and the model with only a single *s* only differed by 2.74, indicating that the model with two different *s* values is not strongly supported.

### Assessment of alternative transmission function forms

We considered several alternative functional forms of the relationship between force of infection and viral titers: linear, threshold, and Hill models. For each of these forms, we continue to assume that viral titers below the LOD do not contribute to the force of infection and thus the probability for transmission. The linear form assumes that viral titers are linearly related to the force of infection, see equation 5.

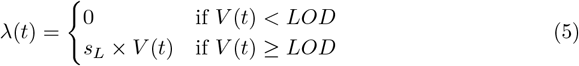

The threshold form assumes that there is some viral titer threshold *h*, below which the force of infection is zero and above which the force of infection is a constant *r*, see equation 6.

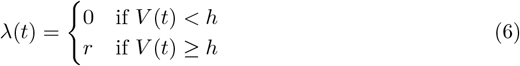

Finally, viral titers could relate to the force of infection according to a Hill function, see equation 7.

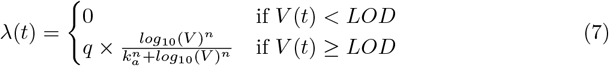

To statistically compare these functional forms, we use a maximum likelihood approach to estimate the required parameters for each form, for both Cal/2009 and Hong Kong/1968. We then use these maximum likelihoods and the number of degrees of freedom in each model to calculate each model’s AICc. Table 1 shows these results. Note that for Hong Kong/1968’s Hill function estimation, we found that above a *k*_*a*_ value of 300, likelihood values continued to increase until we reached the highest value we could consider (a value of 3.8 *×* 10^308^). However, the increase was very slight (*<* 0.01 log-likelihood units over this range). We therefore fixed *k*_*a*_ to 6000 during the inference process and estimated only *q* and *n* for Hong Kong/1968’s Hill function.

### Quantification of parameters at the transmission event

To project the number of Cal/2009 secondary cases, we used the viral titer dynamics of the Cal/2009 infected contact animals, which we considered our population of “theoretical” index Cal/2009 animals. As we had done previously for the index animals when estimating the parameter *s*, we used linear interpolation to obtain continuous viral titer levels for each of these contact animals. We performed 1000 stochastic projections for the Cal/2009 virus. For each stochastic realization, we assumed each animal has an average of *l* one-hour long contacts per day for the duration of their infection. (We used *l* = 15 in Figure 4 based upon previous estimates for the number of daily contacts for children [9]. We explore the impact of assuming different contact rates in Figure 5.) To simulate daily contacts for each animal, we randomly choose 150 timepoints and determine the viral titer in the animal at these timepoints. To incorporate uncertainty in *s*, we randomly draw *s* values and accept them for with a probability equal to their likelihood (shown in Figure 2A) using the Metropolis-Hastings algorithm. We draw until there are 1000 accepted values and set *s* to one of these values in each of the 1000 simulations. With *V* (*t*_*i*,*d*_) denoting the animal’s viral titer for contact number *i* on day *d*, we then calculate the probability of successful transmission at each of these timepoints using equation 4, with *T* = 1*/*24 days.

The total number of secondary cases from a given contact animal is then given by 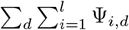 where Ψ_*i,d*_ denotes a random variable that is 1 with probability *Pr*_*i,d*_(+) and 0 otherwise. For each successful transmission (that is, where the random variable Ψ_*i,d*_ is 1), we store the infection time *t*_*i,d*_. This infection time corresponds to the generation interval for this transmission event, defined as the time between infection of an individual and an onward transmission. In Figure 3C, we plot the distribution of these generation intervals for all onward transmissions in the population of Cal/2009 infected contact animals.

We summarize across the 1000 stochastic realizations as follows. We plot in Figure 3E the mean number of secondary infections from the population of Cal/2009 infected contact animals. This mean number corresponds to the basic reproduction number *R*_0_, and we plot the distribution of these *R*_0_ values. To estimate the extent of transmission heterogeneity across these realizations, we follow the approach outlined in [10] and estimate the overdispersion parameter *k* of the negative binomial distribution for each stochastic realization using its simulated distribution of secondary cases. The overdispersion parameter is estimated using the fitdistrplus package in R [23]. We do not estimate *k* for the subset of stochastic realizations that do not result in any secondary cases from any of the animals. In Figure 3F we plot the cumulative distribution function of this overdispersion parameter for the subset of stochastic realizations that had at least one onward transmission. Finally, we calculate the mean generation interval *T*_*c*_ for each stochastic realization and plot the distribution of calculated *T*_*c*_ values across the 1000 simulations in Figure 3G.

Stochastic realizations for Hong Kong/1968 virus are performed in an analogous manner to those for Cal/2009. Summaries across the Hong Kong/1968 stochastic realizations are also performed in an analogous manner. We statistically compare the *R*_0_ and *T*_*c*_ estimates for Cal/2009 and Hong Kong/1968 using Welch’s two sample t-test.

### Prediction of pandemic potential and dynamics

We next estimate *R*_0_ (and the classical 95% confidence interval) across a range of contact rates according to the methods described in the previous section. In Figure 4B, we show the probability of viral establishment across a range of contact rates for two different transmission heterogeneity parameterizations: one in which the overdispersion parameter *k* = *∞* (corresponding to a Poisson distribution of secondary infections across individuals) and one in which the overdispersion parameter *k* = 1 (corresponding to a geometric distribution of secondary infections across individuals). Probabilities of establishment are calculated using the probability generating function:

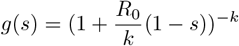

as described in detail in [10].

In Figure 4C, we plot the average length of a stuttering transmission chain. For contact rates that result in an *R*_0_ *<* 1 for a virus, the average length *µ* of a stuttering chain is given by 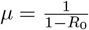.

In Figure 4D, we plot the intrinsic growth rate *r* of a viral epidemic that would establish under a given contact rate. We estimated the intrinsic growth rates under three assumptions for the generation interval distribution [13]. First, we assume the generation interval is exponentially distributed such that 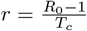. Second, we assume no variance in the generation interval, in which case 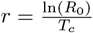. Finally, we assume the generation interval is gamma distributed such that 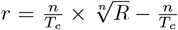. We assume a generation time of 3.6 days because this length is intermediate between the generation times we found for Cal/2009 and Hong Kong/1968.

## Notes

### Competing Interest Statement

The authors have declared no competing interest.

### Summary of Updates

Section "Assessment of alternative transmission function forms" and Fig 3 was added; Fig 4-5 revised. General text updates.

